# IGF-1 Differentially Modulates Glutamate-Induced Toxicity and Stress in Cells of the Neurogliovascular Unit

**DOI:** 10.1101/2021.08.08.455595

**Authors:** Cellas A. Hayes, Brandon G. Ashmore, Akshaya Vijayasankar, Jessica P. Marshall, Nicole M. Ashpole

**Affiliations:** Department of BioMolecular Sciences, University of Mississippi School of Pharmacy, University of Mississippi, University, MS 38677, USA; Research Institute of Pharmaceutical Sciences, University of Mississippi School of Pharmacy, University of Mississippi, University, MS 38677, USA

**Keywords:** Astrocytes, Endothelial Cells, Neurotoxicity, Somatomedin C, Reactive Oxygen Species

## Abstract

The age-related reduction in circulating levels of insulin-like growth factor-1 (IGF-1) is associated with increased risk of stroke and neurodegenerative diseases in advanced age. Numerous reports highlight behavioral and physiological deficits in blood-brain barrier function and neurovascular communication when IGF-1 levels are low. Administration of exogenous IGF-1 reduces the extent of tissue damage and sensorimotor deficits in animal models of ischemic stroke, highlighting the critical role of IGF-1 as a regulator of neurovascular health. The beneficial effects of IGF-1 in the nervous system are often attributed to direct actions on neurons; however, glial cells and the cerebrovasculature are also modulated by IGF-1, and systemic reductions in circulating IGF-1 likely influence the viability and function of the entire neuro-glio-vascular unit. We recently observed that reduced IGF-1 led to impaired glutamate handling in astrocytes. Considering glutamate excitotoxicity is one of the main drivers of neurodegeneration following ischemic stroke, the age-related loss of IGF-1 may also compromise neural function indirectly by altering the function of supporting glia and vasculature. In this study, we assess and compare the effects of IGF-1 signaling on glutamate-induced toxicity and reactive oxygen species (ROS)-produced oxidative stress in primary neuron, astrocyte, and brain microvascular endothelial cell cultures. Our findings verify that neurons are highly susceptible to excitotoxicity, in comparison to astrocytes or endothelial cells, and that a prolonged reduction in IGFR activation increases the extent of toxicity. Moreover, prolonged IGFR inhibition increased the susceptibility of astrocytes to glutamate-induced toxicity and lessened their ability to protect neurons from excitotoxicity. Thus, IGF-1 promotes neuronal survival by acting directly on neurons and indirectly on astrocytes. Despite increased resistance to excitotoxic death, both astrocytes and cerebrovascular endothelial cells exhibit acute increases in glutamate-induced ROS production and mitochondrial dysfunction when IGFR is inhibited at the time of glutamate stimulation. Together these data highlight that each cell type within the neuro-glio-vascular unit differentially responds to stress when IGF-1 signaling was impaired. Therefore, the reductions in circulating IGF-1 observed in advanced age are likely detrimental to the health and function of the entire neuro-glio-vascular unit.

## INTRODUCTION

Aging is a primary risk factor for a number of diseases and pathologies, including stroke, which is recognized as one of the leading causes of death and disability globally (Johnson et al., 2016;Benjamin et al., 2019). Ischemic stroke is particularly devastating as the long-term phenotypic deficits in cognitive behavior and sensorimotor function build in the days following the insult, even if blood flow to the ischemic tissue is restored (as reviewed by (George and Steinberg, 2015)). Ischemia precipitates cellular damage in the brain within minutes, as the loss of blood flow induces metabolic and oxidative stress in the energy-demanding neurons, and nearby glial and cerebrovascular cells. Immediate restoration of blood flow is critical, as the length of ischemia correlates with the extent of tissue damage (Terasaki et al., 2014). To date, the most effective ischemic stroke therapy is fibrinolytic breakdown of the clot with administration of tissue plasminogen activator. Unfortunately, this therapy is time-restricted and requires confirmation of ischemic vs hemorrhagic manifestation, which restricts clinical utility and has limited its use to around 5% of stroke patients each year (National Institute of Neurological and Stroke rt, 1995). Longitudinal observation-based clinical assessments have identified multiple possible biomarkers of stroke risk and severity, including insulin-like growth factor-1 (IGF-1), which may serve as a point of therapeutic intervention in the days surrounding insult (Katzan et al., 2000;2005;Qureshi et al., 2005;Kleindorfer et al., 2008).

IGF-1 is a pleiotropic neuroendocrine modulator that has been assessed in both clinical and preclinical studies of stroke (as reviewed by (Hayes et al., 2021)). IGF-1 levels decline in advanced age, and the extent of this decline is associated with increased risk of stroke incidence, increased mortality, and worsened functional outcomes post-stroke in humans (Johnsen et al., 2004;De Smedt et al., 2011;Tang et al., 2014;Armbrust et al., 2017;Saber et al., 2017). Moreover, altered IGF-1 levels are also associated with increased risk of multiple age-related neuropathologies marked by neurovascular distress, excitotoxicity, and oxidative stress (Watanabe et al., 2005;Sonntag et al., 2013;Ghazi Sherbaf et al., 2018;Gubbi et al., 2018). While many studies associate reduced IGF-1 with increased neurodegeneration, some clinical studies of dementias and stroke see increased IGF-1 levels at the start of pathology, which may be an attempted compensatory mechanism designed to protect the brain (Gubbi et al., 2018;Castilla-Cortazar et al., 2020). Preclinical studies suggest that IGF-1 plays a causal role in the stroke risk and recovery processes by directly regulating cell damage during ischemia (as reviewed by(Hayes et al., 2021)). IGF-1 supplementation in rodent models of middle cerebral artery occlusion and photothrombotic stroke reduces the size of infarcted tissue and the accompanying sensorimotor deficits, even when administered in the days after insult (Liu et al., 2001;Bake et al., 2014;Bake et al., 2016;Parker et al., 2017;Serhan et al., 2019;Hayes et al., 2021). Because IGF-1 regulates numerous cells throughout the body, the cellular mechanism by which IGF-1 protects viability and function in the brain remains unclear. It is commonly inferred that the beneficial effects of IGF-1 are mediated by direct regulation of neurons. However, neurons, astrocytes, endothelial cells, and perivascular cells each express the receptor for IGF-1, IGFR. Together, these cells work to compose the neurovascular unit (also known as neuro-glio-vascular unit) which tightly coordinates physiological responses within the nervous system and is a central target for dysfunction in neurodegenerative diseases and pathologies (Stanimirovic and Friedman, 2012). Additional work is needed to tease apart the impact of each cell type when exogenous IGF-1 is administered to protect against the damage induced by stroke and neurodegenerative diseases.

Animal models of circulating IGF-1 deficiency exhibit impaired neurovascular coupling, compromised blood-brain barrier integrity, and increased prevalence of microhemorrhages (Toth et al., 2015a;Tarantini et al., 2016b;Tarantini et al., 2017;Hayes et al., 2021;Tarantini et al., 2021).

Increased reactive oxygen species (ROS) production, increased inflammation, and reduced glutamate and glucose handling machinery accompany the loss of circulating IGF-1 in these models (Toth et al., 2015b;Toth et al., 2017). Together, these data suggest that the age-related reduction in IGF-1 likely leads to severe consequences on the structure and function of the neuro-glio-vascular unit. Indeed, in vitro studies have highlighted roles for IGF-1 in promoting specific functions of each cellular component of the neuro-glio-vascular unit. Pharmacological and genetic manipulations of IGF-1 alter the viability and oxidative stress levels of cultured neurons (Wang et al., 2014;Li et al., 2017;Chen et al., 2019). Additionally, IGF-1 signaling in brain vascular endothelial cell cultures promotes angiogensis and tight-junction formation/integrity (Lopez-Lopez et al., 2004;Bake et al., 2016;Higashi et al., 2020). We recently reported that deficiency in IGF-1 signaling in astrocytes impairs glutamate handling in vitro and in vivo, by reducing glutamate transporter expression and availability at the cell surface (Prabhu et al., 2019). Considering glutamate overexcitation is a key driver of tissue damage following ischemic stroke (Choi and Rothman, 1990;Moskowitz et al., 2010), it is likely that the age-related loss of IGF-1 may compromise the neuro-glio-vascular unit by weakening the glutamate-buffering capabilities of astrocytes which ultimately exacerbates calcium imbalances, mitochondrial dysfunction, and oxidative stress. While the aforementioned studies highlight the ability of IGF-1 to regulate neurons, astrocytes, and brain-derived endothelial cells, each of these studies was performed under varied conditions. A comparative approach is needed to better-understand how the age-related loss of IGF-1 influences cells within the neuro-glio-vascular unit. Thus, we exposed individual cell cultures and co-cultures to excitotoxic levels of glutamate when IGF-1 signaling was impaired. This approach allows for determination of the cellular mechanism(s) by which IGF-1 exerts its protective effects during ischemic stroke-induced stressors. In addition, it highlights cell-specific responses to a common driver of neuronal dysfunction-glutamate excitotoxicity-which has applications to numerous age-related neurodegenerative disease states.

## MATERIALS AND METHODS

### Animals

All procedures were approved by the Institutional Animal Use and Care Committees of the University of Mississippi, and performed in accordance with their approved guidelines. Timed-pregnant Sprague Dawley rats (16-18 days postcoital plug) were purchased (Envigo) and were temporarily housed until the time of euthanasia in 19 × 11.5 × 11 inch polycarbonate cates. Rats were given access to standard rat chow (Teklad 7001) and water *ad libitum*. At the time of euthanasia, pregnant dams were anesthetized with isofluorane and cervical dislocation was completed. Embryonic pups (E17-19) were excised and rapidly decapitated. Cerebral tissue of both male and female pups were combined for tissue culture isolation.

### Neuron Cultures

Primary rat neuron cultures were established following previously described protocols (Ashpole and Hudmon, 2011). Cell culture dishes and coverslips were coated with Poly-D-Lysine (PDL) (Sigma P6407) for at least 2 hours at 37°C in a humidified incubator containing 5% CO2. Individual cultures were plated in 96-well plates, while 15mm glass coverslips were used to plate neurons for the triple co-cultures. Following PDL coating, cell culture surfaces were washed with 1X PBS (Gibco 10010-023) and dried. Primary neuronal cultures were derived from the cortex and hippocampus of E17-19 Sprague Dawley rat pups. External cerebrovasculature was removed, the tissue was minced and subsequently enzymatically and mechanically digested with papain (Worthington Biochemical Corporation, LS003126) and glass-blown pipettes, respectively. Dissociated cells were pelleted with centrifugation (500-1000g, 5 min) and resuspended in neuron growth media (Neurobasal medium (Gibco 21103-049) containing B27 (Life Technologies, 17504044), penicillin/streptomycin (10 units/mL; Life Technologies, 15140122), and 1x L-glutamine (250303-081)) with a target density of 2,500,000/mL. Partial media changes occurred every 3-4 days post seeding.

### Astrocyte Cultures

Tissue was isolated, digested, and pelleted following the methodology described above for neuron cultures; however, cell pellets were resuspended in astrocyte growth media containing Neurobasal medium (Gibco 21103-049) with 10% fetal bovine serum (Corning 35-010-CV), penicillin/streptomycin (10 units/mL; Life Technologies, 15140122), and 1x L-glutamine (250303-081). Cells were plated on PDL-coated 10cm dishes and media was completely exchanged the day following plating. To select for astrocytes, the cells were subcultured every 3-4 days (90-100% confluent). For this, the growth media was removed, cells were washed with 1X PBS, and 0.05-0.25% of Trypsin-EDTA 1X (Gibco 25200-072) was added. To avoid excess debris, trypsin was applied twice during the first passage-once for 2 minutes to dislodge neurons and microglial cells from the upper layer (vacuumed off), and a second time to release the astrocytes underneath. Once the astrocytes dislodged, the cells were centrifuged (1000xg; 5 min) and resuspended in growth medium for further plating.

### Endothelial Cell Culture

Rat brain microvascular endothelial cells (RBMVECs) and medium were purchased from Cell Applications Inc. (San Diego) and cultured following the manufacturer’s guidelines. In brief, cells were grown on a T-75 flasks pre-coated with Cell Attachment Factor Solution (123-100, Cell Applications), in 15mL of RBMVEC growth medium (R819-500). Growth medium was changed 24 hours after initial seeding, and cells were sub-cultured every 4-6 days or when at 80-100% confluency. Media was changed every other day. RBMVECs were passaged using the same protocol as astrocytes, using the proprietary trypsin solution (Cell Applications Inc).

### Co-cultures

Triple cultures were developed by plating primary neurons on glass coverslips, RBMVECs on transwell inserts, and astrocytes on the surface of a multi-well plate. Coverslips were elevated off the surface of the multi-well plate and astrocytes were seated in the outer 1/3 of the wells. The media on all 3 cell types was exchanged for neuron growth media 24 hours prior to combination of the co-cultures to allow for uniformity between media types.

### Cells treatments

Glutamate stimulation was performed using L-glutamic acid and glycine (FisherScientific) in a ratio of 10:1 glutamate:glycine diluted from a PBS stock solution. Increasing concentrations of glutamate were prepared in the appropriate cell growth media, and administered as a 2x solution to the cells plated in their respective media (final concentration 1x upon application). Cells were incubated for one hour at 37°C. Following treatment, glutamate media was removed, cells were washed in fresh growth media twice, and incubated in new growth media until the designated time point. All cell viability studies were conducted twenty-four hours after cells were stimulated with glutamate. All other study timepoints are described in the respective sections. In one experimental series, media lacking the B-27 supplement was administered 24 hours prior to glutamate stimulation.

Stock solutions of picropodophyllin toxin (PPP; Sigma T-9576) and MK-801 (Sigma M107) were prepared in dimethyl sulfoxide (DMSO; Fisher BP321-100) and administered to cells at concentrations and time-points described. Treatment groups were block randomized to ensure equal sampling.

### Cell viability

Following treatments, cell viability/cytotoxicity was quantified using Live:Dead viability/cytotoxicity kit (Thermo Fisher Scientific L3224). Calcien-AM and ethidium homodimer were diluted (1:1000-2000; 1:500-1000) were diluted in PBS. Growth media was removed and cells were washed with 1X PBS prior to adding the live/dead labels. After 20 minutes of incubation, cells were imaged on the Nikon Ti2-E HCA inverted fluorescent microscope using the JOBS automated image acquisition (Nikon). Magnification was set to 200x (20x, extra-long working distance objective) and samples were excited with the LED Triggered acquisition exposures using excitation/emission filters for 470-FITC and 540-TRITC. The JOBS program selected 6 fields at random per well for imaging, and the total number of live and dead cells were quantified in 2-3 images using the Nikon Elements Cell Analysis plug-in features. The total number of cells and percent toxicity were calculated by observers blinded to treatment groups.

### ROS quantification

Following treatment, reactive oxygen species (ROS) production was quantified in astrocytes and endothelial cells. Media was removed, and cells were washed with 1X PBS. Cells were then loaded with 3μM of 2’,7’-dichlorodihydrofluorescein diacetate (DCFDA) (Thermofisher, D399) for 20 minutes and incubated at 37°C in a humidified incubator containing 5% CO^2^. DCFDA was then imaged using the FITC filter on the Nikon Ti2-E microscope as described above. Six images at random through the JOBS interface and 4-6 non-adjacent cells per field were selected at random to be quantified by a blinded observer. DCFDA intensity of each cell was normalized to the background of that image.

### Seahorse

Cells were seeded on the XFe96 growth plates 2-3 days prior to extracellular efflux assessment. Treatments were administered at the desired timepoints and mitochondrial response was assessed using the Cell Mito Stress Test, per manufacturer recommendations. The sensor cartridge was hydrated in water overnight at 37°C and then equilibrated with the recommended equilibration buffer. Oligomycin (1.5μM), FCCP (1μM), and rotenone (0.5μM) were loaded into the sensor plates and injected into the wells by the XFe/XF Seahorse system. Basal and maximal respiration, as well as proton leak were auto-calculated by the Seahorse program using an average of 3 readings per stage.

### Data Analysis

Statistical analysis and graphical analyses were performed using Sigma Plot version 14 software and R studio. One-way and two-way ANOVAs was used when appropriate (defined in the figure legend). Post-hoc comparisons were selected based upon the experimental question and fulfillment of normality and equal distribution. Details for each are provided in figure legend. All data were expressed with mean +/− standard error. Sample sizes were estimated using previous variance observations with these experimental methodologies. With β=0.8, n=4-6 was required to achieve power for most studies. F values for two-way ANOVAs are described in the results and main effects are noted in figures with a pound sign (#). For all studies, p<0.05 was used as the statistical significance value and denoted using an asterisk (*).

## RESULTS

### IGF-1 protects against excitotoxicity in neurons

To determine if inhibition of IGF-1 signaling differentially affects the glutamate sensitivity of cells within the neuro-glio-vascular unit, primary cultures of neurons, astrocytes, and microvascular endothelial cells were established. Neurons are sensitive to excitotoxicity when exposed to high concentrations of glutamate and glycine, due to overactivation of ionotropic glutamate receptors (GluNR). As expected, acute treatment of neurons with 100μM glutamate/10μM glycine (termed glutamate stimulation throughout) resulted in a significant increase in cellular toxicity 24 hours after stimulation (Figure 1A-B; p<0.001 vs unstimulated). Co-administration of the GluNR antagonist MK-801 prevented excitotoxicity (Figure 1B; p<0.001 vs glutamate stimulated control). Administration of exogenous 100nM IGF-1 at the time of stimulation also prevented excitotoxicity (p=0.019 vs glutamate stimulated control), consistent with previous reports. No differences in the total number of neurons per field were noted following treatment (Supplemental Figure 1).

**Figure 1:**
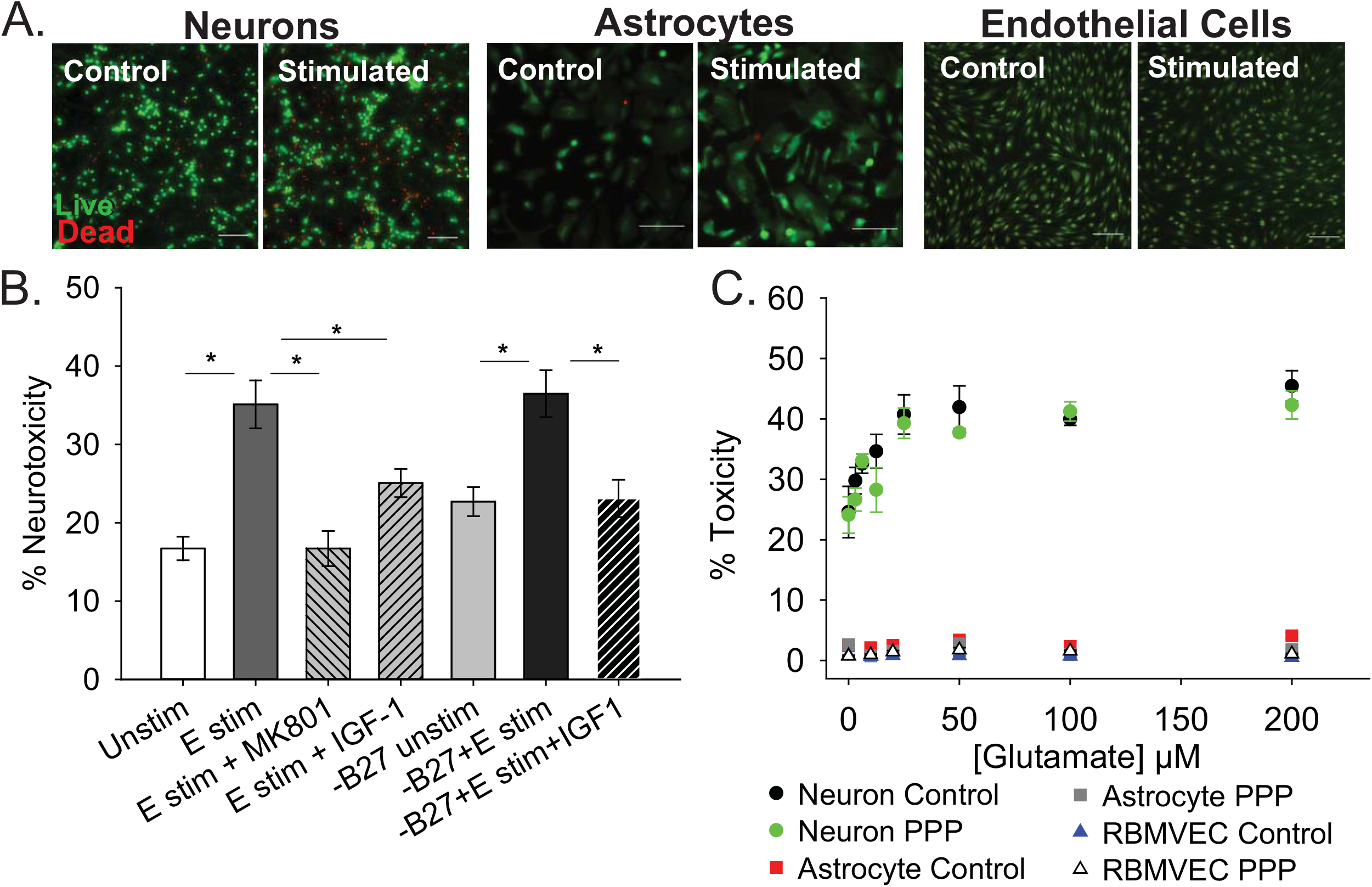
IGF-1 protects against excitotoxicity in glutamate-sensitive neurons. (A) Representative images of neurons (left two panels), astrocytes (middle two panels), and rat brain microvascular endothelial cells (right two panels) stained with live (green)/Dead (red) viability/cytotoxicity indicator dyes 24 hours after stimulation with control or 100μM glutamate, as indicated. Scale bar represents 100μm. (B) Average toxicity in neurons 24 hours after treatment (n=8 wells/group). A one-way ANOVA with post-hoc Bonferroni was used for statistical comparisons; * indicates significant difference between the delineated bars. (C) Average toxicity of cells 24 hours after acute treatment with both 0.5μM PPP and increasing concentrations of glutamate (n=4-8 wells/group). A two-way ANOVA was employed with glutamate concentrations and PPP-treatment serving as factors. No significant differences were observed. Graph color code: black circles=vehicle-treated neurons; green circles= PPP-treated neurons; red boxes= vehicle-treated astrocytes; dark gray boxes= PPP-treated astrocytes; blue triangles= vehicle-treated rat brain microvascular endothelial cells (RBMVEC); light gray triangles= PPP-treated RBMVEC. All data are presented as mean ± SEM.

The neuronal growth media contains a supplement (B27) that has a high concentration of insulin, which can cross-talk and activate IGFR. Thus, to better determine the protective effects of IGF-1 signaling, the excitotoxicity profile was again assessed without B27 present. No change in baseline levels of toxicity were observed between control neurons with and without B27, and glutamate stimulation still significantly increased toxicity in the absence of B27 (Figure 1B; p=0.002 vs -B27 unstimulated control). Once more, exogenous IGF-1 prevented excitotoxicity (Figure 1B; p=0.003 vs -B27 glutamate stimulated control).

To more specifically assess the impact of reducing IGF-1 signaling, the extent of excitotoxicity when IGFR was pharmacologically inhibited was then examined. For this, picopodophyllinotoxin (PPP), a known small molecule inhibitor of the IGFR receptor kinase activity, was co-administered at the time of glutamate stimulation and toxicity was measured 24 hours later. Surprisingly, PPP did not shift the concentration-dependence or maximal extent of glutamate toxicity in cultured neurons (Figure 1C). A two-way ANOVA revealed a significant effect of glutamate concentration (F=13.759, p<0.001), but no significant effect of PPP treatment (F=2.598, p=0.111), and no significant interaction between the two factors (F=0.506, p=0.827).

Astrocytes and microvascular endothelial cells are not as sensitive to glutamate as neurons as no change in toxicity was observed across all glutamate concentrations (Figure 1A-C). Similarly, no increase in sensitivity was observed with IGFR inhibition at the time of glutamate stimulation (Figure 1C).

### Prolonged IGFR inhibition increases astrocytic sensitivity to glutamate

The lack of effect with pharmacological inhibition of IGFR in Figure 1C was inconsistent with the protective effects of exogenous IGF-1 in Figure 1B. However, co-administration of PPP at the time of glutamate could have temporal confounds that limit interpretation. Thus, we next administered PPP 24 hours prior to glutamate stimulation to better mimic the prolonged loss of IGFR signaling observed in aging. Prolonged IGFR inhibition with 0.5μM PPP did not lead to difference in neuronal excitotoxicity (F=0.255, p=0.615), but did lead to differences in astrocyte viability (F=6.677, p=0.011) (Figure 2A-B). Within group comparisons did not unveil a specific concentration of glutamate that PPP treatment significantly worsened, thus it is assumed that IGFR inhibition led to modest reductions in cell viability across all glutamate treatments.

**Figure 2:**
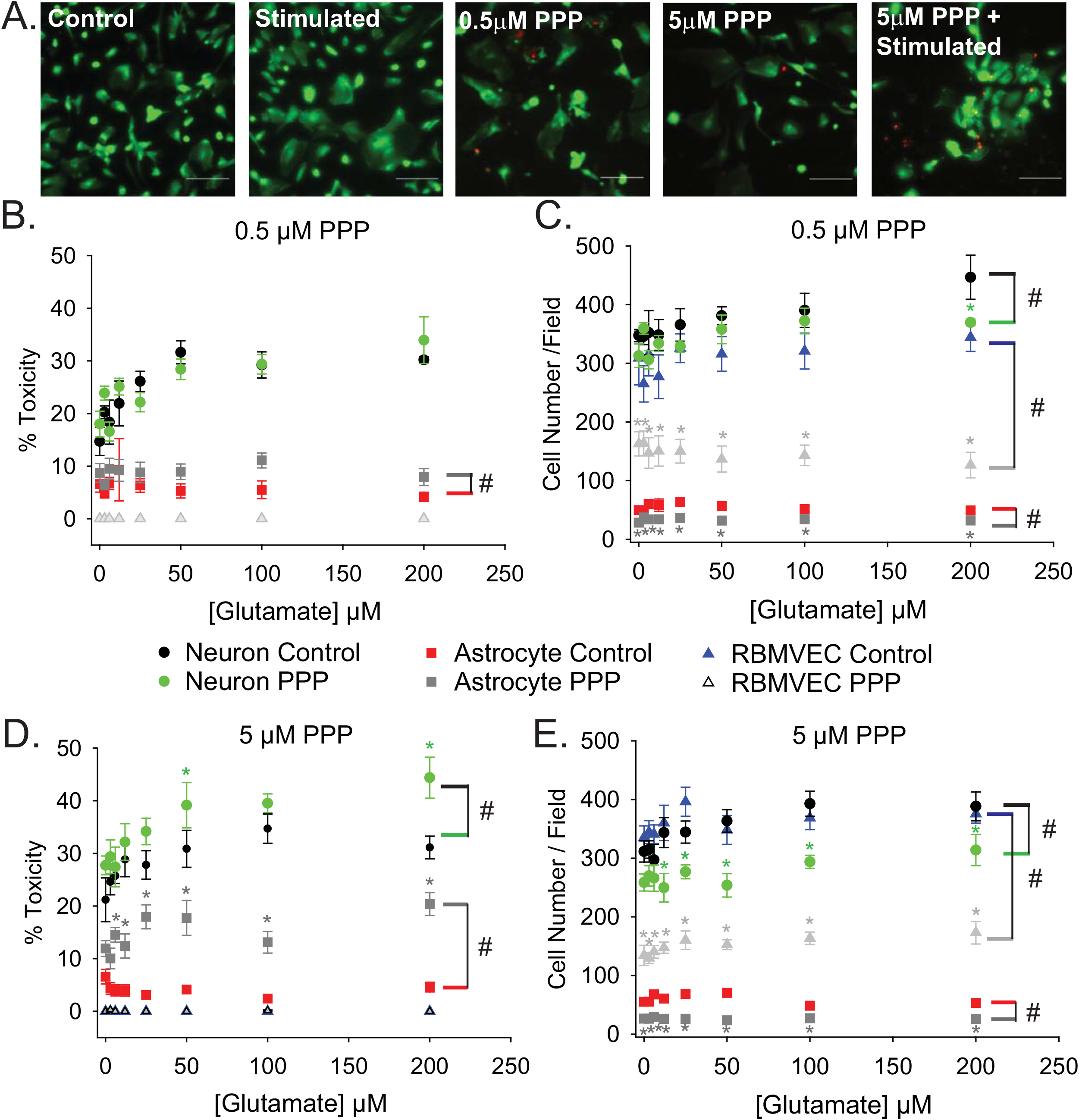
Prolonged IGFR inhibition increases astrocytic sensitivity to glutamate and reduces astrocyte and endothelial cell number. (A) Representative images of astrocytes stained with live (green)/Dead (red) viability/cytotoxicity indicator dyes. From left to right: unstimulated control, glutamate-stimulated vehicle-treated, unstimulated 0.5μM PPP-treated, unstimulated 5μM PPP, and glutamate-stimulated 5μM PPP. Scale bar represents 100μm. (B) Average toxicity and (C) average cell number per well of cells pre-treated with 0.5μM PPP, and subsequently stimulated with increasing concentrations of glutamate, 24 hours after stimulation. (D) Average toxicity and (E) average cell number per well of cells pre-treated with 5μM PPP, and subsequently stimulated with increasing concentrations of glutamate, 24 hours after stimulation. All data are presented as mean ± SEM, n=5-8 wells/treatment group. A two-way ANOVA was employed with glutamate concentrations and PPP-treatment serving as factors. Cell types were each analyzed independently. # indicates a significant main effect within that cell type. Post-hoc Holm-Sidak tests were used for pairwise multiple comparisons. *indicates a significant difference between PPP-treatment and vehicle-treatment within a given glutamate concentration. Graph color code: black circles=vehicle-treated neurons; green circles= PPP-treated neurons; red boxes= vehicle-treated astrocytes; dark gray boxes= PPP-treated astrocytes; blue triangles= vehicle-treated rat brain microvascular endothelial cells (RBMVEC); light gray triangles= PPP-treated RBMVEC.

No changes in endothelial cell viability were observed with prolonged IGFR inhibition; however, a pronounced reduction in total cell number was noted when microvascular endothelial cells were treated with 0.5μM PPP (F=117.942, p<0.001) (Figure 2C). A significant difference within unstimulated controls with PPP pre-treatment was detected (p<0.001), and this effect persisted across all glutamate concentrations. Thus, no main effect of, or interaction with, glutamate concentration was observed in the treated microvascular endothelial cells.

The change in endothelial cell number prompted further analyses of neuron and astrocyte counts with treatment. Similar reductions in astrocyte count were observed with 0.5μM PPP treatment (F=73.322, p<0.001) (Figure 2A-C). Glutamate concentration did not alter astrocyte count nor interact with the PPP effect. Main effects of both glutamate concentration and PPP treatment on total neuron count were detected (F=3.309, p=0.004; F=7.835, p=0.006, respectively). Post-hoc comparisons within groups revealed that the only significant difference within the PPP vs vehicle treatments was the drop in total cell number at 200 μM glutamate (p=0.012), likely an indication of dead cells washed away during the staining process at this high concentration of glutamate.

When the concentration of PPP was increased to 5 μM, significant differences in neuron and astrocyte survival were observed (Figure 2D). Both glutamate concentration and PPP treatment had significant main effects on neurotoxicity (F=4.272, p<0.001; F=12.565, p<0.001, respectively). However, no interaction between the two factors was detected (F=0.732; p=0.646). Within group comparisons revealed significant increases in neurotoxicity when neurons pretreated with 5 μM PPP were stimulated with 50 and 200 μM glutamate (p=0.048 and p=0.006, respectively). No baseline differences in toxicity amongst unstimulated controls treated with vehicle or PPP were observed (p=0.751). A significant reduction in total neuron number was observed with 5 μM PPP treatment and with glutamate (F=49.058, p<0.001; F=2.878, p=0.009, respectively) (Figure 2E). No significant interactions between the factors was detected (F=0.932, p=0.485). Within factor comparisons showed decreased cell number within PPP-treated neuron cultures stimulated with ≥12.5μM glutamate, again suggesting potential loss of dead cells during the staining process since neurons are non-mitotic.

Prolonged IGFR inhibition in astrocytes also resulted in main effects of glutamate concentration and PPP treatment on the levels of cytotoxicity (F=157.321, p<0.001; F=2.209, p=0.039, respectively) (Figure 2D). Moreover, a significant interaction between glutamate concentration and PPP treatment was also observed (F=2.743, p=0.011). Within factor comparisons revealed significant increases in PPP-pretreated astrocytes stimulated with ≥6μM glutamate, suggesting that while astrocytes display increased resistance to glutamate toxicity under control conditions, the loss of IGF-1 signaling predisposes astrocytes to excitotoxic stress. A significant reduction in astrocyte cell number was also observed with 5 μM PPP treatment (F=257.893, p<0.001), across all glutamate concentrations. Main effect differences for glutamate concentration did not reach significance (F=1.882, p=0.079), nor did interactions between factors (F=2.071, p=0.052).

Similar to the before no differences in endothelial cell toxicity were observed across any treatment groups, but 5 μM PPP did reduce cell number across all glutamate concentrations (F=538.084, p<0.001) (Figure 2E). Again, this suggests that while IGFR inhibition did not predispose endothelial cells to glutamate toxicity, it likely reduces cell division.

### Astrocytes fail to protect neurons when IGFR is inhibited

Cells within the neuro-glio-vascular unit coordinate to create an optimal microenvironment, by releasing growth mediators, buffering stressors, and regulating nutrient supply. Thus, we next assessed whether co-cultures of endothelial cells, neurons, and astrocytes would exhibit differences in glutamate sensitivity when IGFR is inhibited. As anticipated, 100 μM glutamate did not significantly increase neurotoxicity in the presence of astrocytes and endothelial cells, nor did acute co-administration of 0.5 μM PPP with 100 μM glutamate (p=0.139) (Figure 3A). No differences in astrocyte or endothelial cell toxicity were observed in these acute treatments.

**Figure 3:**
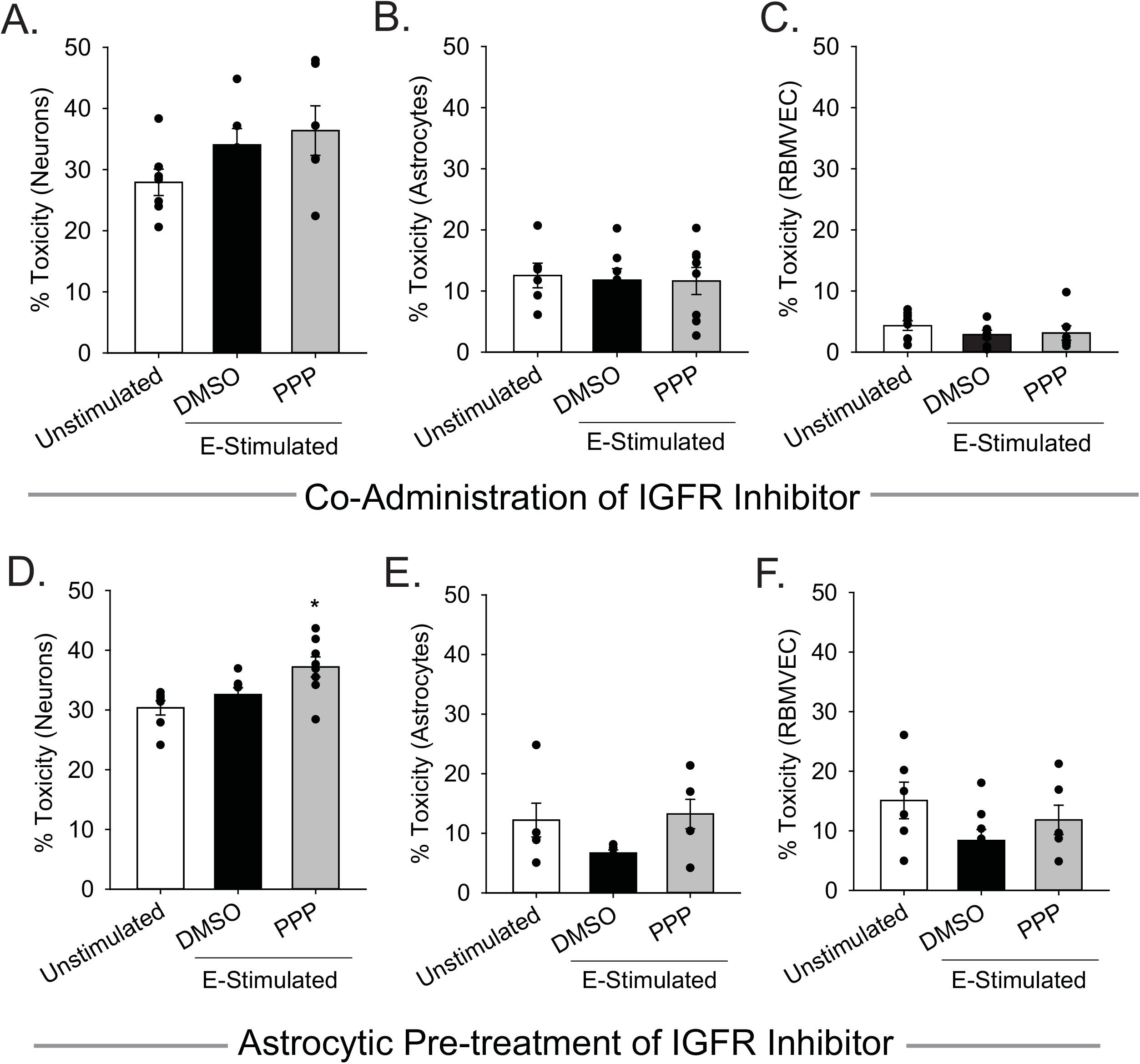
Astrocytes fail to protect against excitotoxicity when IGFR is inhibited. Average cell death 24 hours after triple cultures of neurons (A), astrocytes (B), and endothelial cells (C) were co-administered 0.5μM PPP and 100μM glutamate for one hour (n=6-8 wells/group). Oneway ANOVA failed to detect differences in any of the treatment groups. (D-F) Astrocytes were pre-treated with 5μM PPP for 24 hours prior to combining with endothelial cells and neurons for a triple culture system. Average cell death in the neurons (D), astrocytes (E), and endothelial cells (F) after stimulation with 100μM glutamate for one hour (n=5-8 wells/group). One-way ANOVAs with post-hoc Bonferroni comparison was used for statistical analyses. *indicates significant difference between control and PPP-treated glutamate-stimulated neurons. All data are presented as mean ± SEM, n=5-8 wells/treatment group.

When astrocytes alone were pre-treated with 5 μM PPP and then combined with neurons and endothelial cells for glutamate stimulation, a significant increase in neurotoxicity was observed (p=0.007 vs control). No differences in toxicity or cell number were noted in either the astrocytes or endothelial cells within the triple cultures (Supplemental Figure1). Together these data suggest that a reduction of IGFR signaling in astrocytes impairs their ability to buffer glutamate and protect neurons from overexcitation, even when in the presence of endothelial cells.

### Astrocytic and endothelial ROS levels are elevated by IGFR inhibition and glutamate stimulation

While acute co-administration of 0.5μM PPP and glutamate did not increase toxicity in astrocyte or endothelial cultures, alterations in metabolic function and oxidative stress may still occur with this stressor. Thus, astrocytes and endothelial cells were treated with 100μM glutamate alone, or in conjunction with 0.5μM PPP, and the production of reactive oxygen species (ROS) was quantified at various time points following treatment. Astrocytic ROS production increases in the hours following glutamate stimulation (Figure 4A). Significant effects of both time and treatment were observed (F=185.57, p<0.001; F=3.187, p=0.027, respectively), as well as an interaction between both factors (F=4.586, p<0.001). Five hours post-treatment, vehicle treated and PPP-treated astrocytes stimulated with glutamate showed significant increases in ROS levels over unstimulated controls (both p<0.001 vs unstimulated control). Moreover, co-administration of PPP and glutamate resulted in significantly increased ROS levels than glutamate stimulated controls (p=0.014), suggesting that the extent of glutamate-induced ROS production was exacerbated by IGFR inhibition. Additional analysis of mitochondrial bioenergetics at this time point revealed that glutamate stimulation decreased maximal extracellular acidification rates, while PPP+glutamate decreased extracellular acidification and oxygen consumption rates, suggesting changes in both glycolysis and oxidative phosphorylation (Supplemental Figure 2). No differences in basal respiration rates were noted with any of the treatments (Supplemental Figure 2).

**Figure 4:**
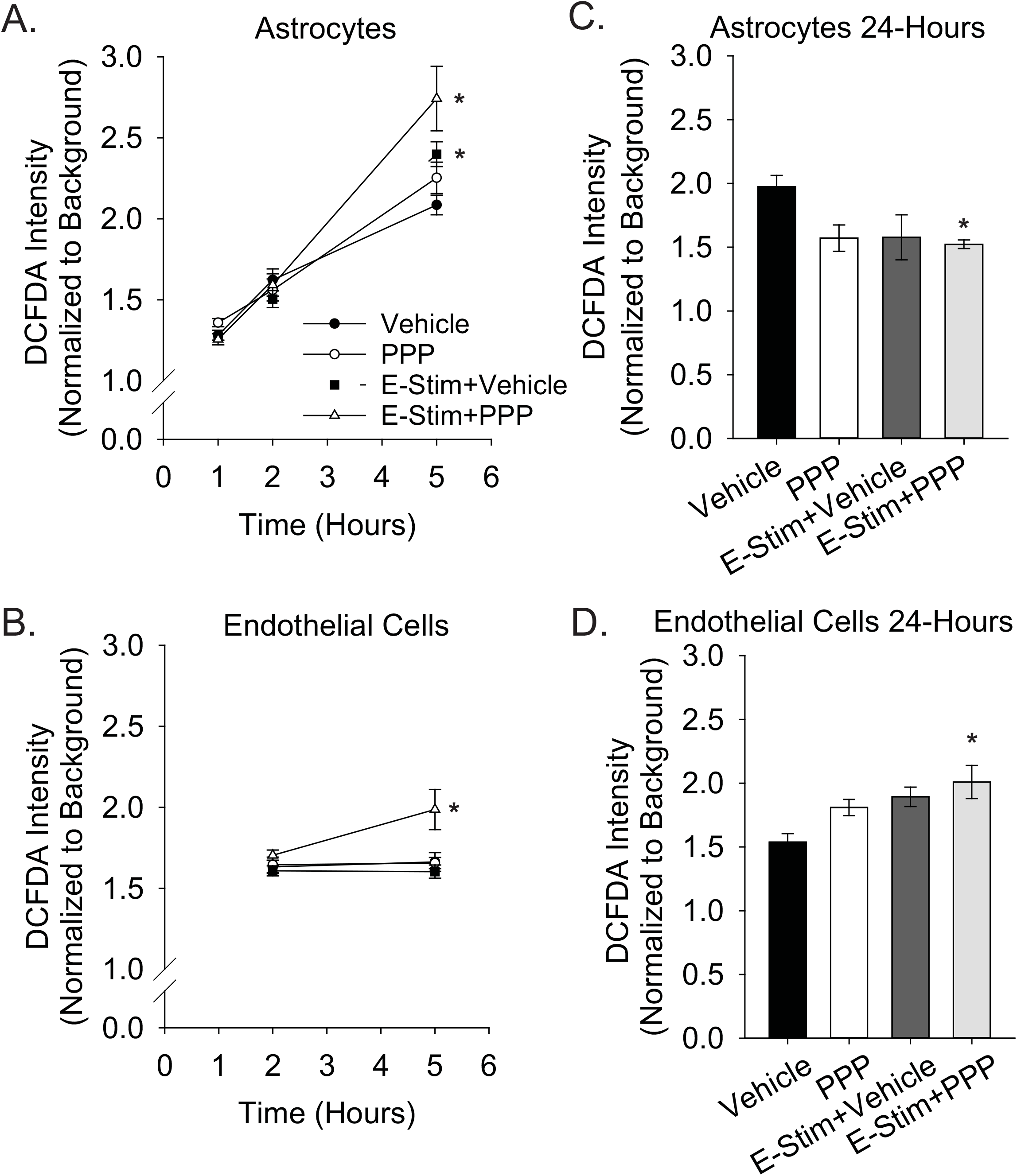
Astrocytic and endothelial ROS levels are elevated by IGFR inhibition and glutamate stimulation. Average ROS levels in pure cultures of astrocytes (A) or endothelial cells (B) co-treated with 0.5μM PPP and 100μM glutamate for one hour, and subsequently measured one hour, two hours, and five hours after treatment. Four cells were selected at random per field, with 3 fields per well, and 8-10 wells per treatment group. Background intensity within each image was measured and used to normalize. A two-way ANOVA was employed with both time and treatment as factors, and a post-hoc Holm-Sidak was used to compare within groups. * indicates a significant effect of treatment at that time. Average ROS levels in pure cultures of astrocytes (C) or endothelial cells (D) co-treated with 0.5μM PPP and 100μM glutamate for one hour, and subsequently measured 24 hours after treatment. Four cells were selected at random per field, with 3 fields per well, and 4-8 wells per treatment group. A one-way ANOVA with post-hoc Dunnett’s test was used for statistical comparisons to control. * indicates a significant effect vs control. All data are presented as mean ± SEM

Similar to astrocytes, brain-derived microvascular endothelial cells showed increased ROS production five hours after 0.5μM PPP was co-administered with 100μM glutamate. A main effect treatment was observed (F=5.287, p=0.004). While glutamate alone did not increase ROS production in the endothelial cells (p=0.945), a significant increase in ROS was seen in cells treated with PPP+glutamate versus unstimulated control (p=0.02), and versus glutamate-stimulated control (p=0.004).

Additional analyses of ROS levels twenty-four hours following treatment revealed differential recovery from acute IGFR inhibition and glutamate stimulation. Astrocytes do not show increased ROS at 24 hours with 100μM glutamate or 100μM glutamate+0.5μM PPP treatment (Figure 4C). In fact, ROS levels were significantly reduced in glutamate-stimulated PPP-treated astrocytes vs unstimulated controls (p=0.024) and PPP treatment was trending (p=0.053 vs control). Endothelial cells continue to show increased ROS levels long after the acute exposure to 100μM glutamate+0.5μM PPP (p=0.013 vs unstimulated control) (Figure 4D), suggesting that IGFR inhibition at the time of glutamate stimulation increases endothelial ROS levels long-term. Note, this was with acute PPP treatment, at a time point in which no differences in endothelial cell toxicity was observed (Figure 1C).

## DISCUSSION

The age-related loss of IGF-1 has been linked to cognitive impairment, neurodegeneration, and increased susceptibility ischemic stroke and other neurovascular pathologies (Sonntag et al., 2013;Ashpole et al., 2015b). These deficits often come as a disconnect from the enhanced longevity observed in animal models of IGF-1 deficiency. While early-life reductions in circulating IGF-1 do lead to increased lifespan, reduced IGF-1 in advanced age has multiple consequences in animal models as it increases sarcopenia, bone frailty, learning and memory deficits, and cerebrovascular dysfunction (Ashpole et al., 2015a;Toth et al., 2015b;Ashpole et al., 2016;Tarantini et al., 2016a;Ashpole et al., 2017;Farias Quipildor et al., 2019;Tarantini et al., 2021). For example, IGF-1 deficiency in mice decreases stimulation-evoked cerebral blood flow, blood brain barrier integrity, and the strength and flexibility of cerebral arteries. IGF-1 deficiency also impairs the production and release of vasomediator eicosanoids from astrocytes, alters endothelial nitric oxide production, and increases susceptibility to hypertension-induced microhemorrhages (Toth et al., 2015b;Tarantini et al., 2017;Fulop et al., 2019;Tarantini et al., 2021). Considering this, it is not surprising that IGF-1 has long-been implicated in the risk and severity of ischemic stroke. As described earlier, clinical and preclinical evidence highlight inverse correlation (and causation in animal models) between IGF-1 and the risk/outcome of ischemic stroke (Hayes et al., 2021).

While recent studies are beginning to shed light on the impact of IGF-1 on neurovascular communication in advanced age, there remains a significant gap in our understanding of how individual cell types that normally coordinate signaling together within the neurovascular unit each respond to reduced IGF-1 signaling. In this study, we aimed to compare the effects of IGF-1 deficiency on the cellular responses to high levels of extracellular glutamate. As the predominant excitatory neurotransmitter in the brain, glutamate induces depolarization and initiates multiple calcium signaling cascades. During periods of ischemia or proteotoxic stress, neurons experience energy imbalances and ionic stressors that ultimately lead to terminal depolarization and the dumping of glutamate stores, inflammatory mediators, and oxidative stressors into the extracellular space, which expands the region of distress and malfunction (as recently reviewed (Choi, 2020)). This excitotoxic cascade is one of the drivers of neuronal loss and glial activation in ischemic stroke and neurodegenerative diseases. Thus, understanding how the loss of IGF-1 alters the glutamate response of neurons, astrocytes, and endothelial cells within the neurovascular unit provides much-needed information on potential cellular mechanisms by which IGF-1 deficiency in advanced age influences health and function of the brain.

Recent evidence from our laboratory indicated that IGFR inhibition reduced the ability of astrocytes to buffer extracellular glutamate by decreasing glutamate transporter availability (Prabhu et al., 2019). Therefore, in this study, we hypothesized that maintenance of IGF-1 signaling in astrocytes is necessary for neuroprotection from glutamate excitotoxicity. We also hypothesized that maintenance of IGF-1 signaling in neurons was essential for limiting overexcitation, as other studies have shown exogenous IGF-1 can protect against excitotoxicity and oxidative stress (Wang et al., 2014;Li et al., 2017;Chen et al., 2019). Our results indicate that while exogenous IGF-1 can protect neurons from glutamate, acute and prolonged loss of IGFR signaling does not exacerbate neuronal death in cultures of pure neurons. Interestingly, inhibition of IGF-1R not only impairs the neuroprotective capabilities of astrocytes, it predisposes astrocytes to glutamate toxicity as well. It is unclear what the mechanism for this enhanced sensitivity may be. Perhaps the lack of glutamate transporter availability with IGF-1 signaling deficiency resulted in increased glutamate receptor activation in the astrocytes as well, since astrocytes are known to express a variety of ionotropic and metabotropic glutamate receptors (Serrano et al., 2008;Lalo et al., 2011;Bradley and Challiss, 2012;Ceprian and Fulton, 2019). If this were the case, the astrocytes could be experiencing intracellular calcium imbalances and oxidative stress, which have both been shown to increase astrotoxicity in response to other stressors.

As mentioned, reductions in IGF-1 in the brain increase ROS levels and mitochondrial dysfunction. We observed significant increases in ROS production in both astrocytes and endothelial cells when IGFR was inhibited at the time of glutamate stimulation. We chose to focus our experimental design on these two cell types and this acute treatment because it did not lead to any changes in observed toxicity. Oxidative stress is expected in neurons that are dying from excitotoxicity, but neither astrocytes nor endothelial cells were sensitive to glutamate when IGFR was acutely inhibited. The increased ROS levels in both cell types indicates that the cells were indeed undergoing a stress response with the combined treatment. While it could be inferred that the ability of astrocytes to return ROS levels back to baseline by twenty-four hours may contribute to their increased resistance to excitotoxic stress, there is a disconnect in the endothelial cells that likely renders this to be more complicated. Perhaps an increase in endothelial cell toxicity would have been observed at later time points. Further studies on the underlying mechanism(s) and consequence(s) of these observations are needed.

Combined triple cultures of neurons, astrocytes, and endothelial cells highlighted a few distinct responses to glutamate and IGFR inhibition than the individual cultures of each cell type. First, the same concentration of glutamate that led to significant increases in neurotoxicity within pure neuron cultures failed to induce toxicity within the triple culture. This was an expected outcome, as astrocytes are known to buffer glutamate and co-cultures of neurons and astrocytes have been shown to reduce the extent of neurotoxicity (Choi et al., 1987;Rosenberg et al., 1992). Second, and more interestingly, prolonged IGFR inhibition in astrocytes prior to glutamate stimulation did not increase the toxicity of astrocytes nor decrease the total cell number of astrocytes within the triple cultures. These effects were pronounced in pure astrocyte cultures, suggesting that coculturing astrocytes with neurons and endothelial cells afforded them protection against the stressors of IGFR deficiency/glutamate stimulation. Additional studies examining ROS levels, inflammatory mediators, and growth factor production and release with triple cultures exposed to these same stressors may be warranted.

In our recent study examining glutamate uptake in astrocytes, we utilized similar concentrations of the pharmacological inhibitor PPP and did not observe a significant reduction in total cell number. Thus, we were surprised to see a reduction in astrocyte count here. However, in that study, cells were treated when at or close to confluence, which was evident in DAPI-stained microscopy images accompanying those data (Prabhu et al., 2019). In the current study, we aimed to treat at 75% confluence in order to avoid stress of over-growth, however microscopic live/dead staining analysis showed that even the vehicle-treated controls were not at this level of confluence even 24 hours after treatment. Thus, the reduction in cell number observed in pure cultures of astrocytes and endothelial cells was likely due to changes in cell division and growth with IGF-1 deficiency. Nevertheless, it is interesting that the reduction in astrocyte number with prolonged IGFR inhibition was restricted to the pure cultures, and not the triple cultures.

Our comparative study is limited by not including pericytes, as they are another important cell type within the neurovascular unit. Pericytes not only serve as a protective layer of the blood brain barrier, they regulate blood capillary diameter within the brain by dilating and relaxing in response to glutamate (Kisler et al., 2017;Brown et al., 2019;Kisler et al., 2020). Because of this, pericytes are important during ischemic reperfusion as they can assist with restoring flow of oxygen and glucose to the ischemic tissue (Yang et al., 2017;Khennouf et al., 2018;Sun et al., 2020). Little is known about the influence of IGF-1 on pericytes within the brain, and additional studies on how IGF-1 may influence pericytic glutamate response are needed.

Together, our study highlights that cell types within the neurovascular unit differentially respond to IGF-1 signaling. Despite differences in baseline susceptibility to glutamate, both neurons and astrocytes are both protected from excitoxicity by IGF-1. Growth and division of endothelial cells and astrocytes are both influenced by IGF-1, however endothelial cells remain resistant to glutamate toxicity even when IGF-1 signaling is reduced. The resistance of these supporting cells does not mean there are no consequences of reduced IGF-1 signaling in the time of glutamate stress, as both astrocytes and endothelial cells show signs of oxidative stress in the hours following exposure. The combination of neuron, astrocyte, and endothelial cells in culture afforded astrocytic protection from IGF-1 deficiency/glutamate stress but also highlighted a failure in the ability of astrocytes to protect the nearby neurons. Thus, the age-related loss of IGF-1 likely impairs the function and vitality of the entire neurovascular unit by differentially exerting stressors on neurons, endothelial cells, and astrocytes.

## Author Contributions

CH and NA were responsible for conceptualization, investigation, project administration, and formal statistical analyses. CH, JM, and NA developed and designed methodology. BA, AV, and NA performed blinded image analyses and curated data. CH prepared visualizations. CH, BA, and NA prepared the original draft, and all authors reviewed and edited the manuscript.

## Data Availability Statement

All original contributions presented in the study are available upon request.

---

**Supplemental Figure 1: Additional assessments of total cell number.** (A) Average number of neurons per field 24 hours after treatment. One-way ANOVA revealed no difference between means. (B-D) Astrocytes were pre-treated with 5μM PPP for 24 hours prior to combining with endothelial cells and neurons for a triple culture system. Average number of cells per field in the neurons (B), astrocytes (C), and endothelial cells (D) cultures after stimulation with 100μM glutamate for one hour (n=5-8 wells/group). One-way ANOVA revealed no difference between means, and the p value of the ANOVA is listed for each. All data are presented as mean ± SEM.

**Supplemental Figure 2: Mitochondrial stress assessment in astrocytes.** Average oxygen comsumption rate (y axis) and extracellular acidification rate (x axis) in astrocytes treated with vehicle control, 100μM glutamate, or 100μM glutamate+0.5μM PPP for one hor. Measurements occurred 5 hours after treatment. Maximal respiration is grouped in the top right 3 points, and basal respiration is grouped in the bottom left 3 points. A one-way ANOVA was used for each comparison of maximal OCR, maximal ECAR, basal OCR, and basal ECAR. Post-hoc Dunnett’s test vs vehicle control was used when relevant. * indicates significant difference in OCR, and # indicates significant difference in ECAR. All data are presented as mean ± SEM, n=7-8 wells/group.

## References

(2005). Prioritizing interventions to improve rates of thrombolysis for ischemic stroke. Neurology 64, 654–659.

Armbrust, M., Worthmann, H., Dengler, R., Schumacher, H., Lichtinghagen, R., Eschenfelder, C.C., Endres, M., and Ebinger, M. (2017). Circulating Insulin-like Growth Factor-1 and Insulin-like Growth Factor Binding Protein-3 predict Three-months Outcome after Ischemic Stroke. Exp Clin Endocrinol Diabetes 125, 485–491.

Ashpole, N.M., Herron, J.C., Estep, P.N., Logan, S., Hodges, E.L., Yabluchanskiy, A., Humphrey, M.B., and Sonntag, W.E. (2016). Differential effects of IGF-1 deficiency during the life span on structural and biomechanical properties in the tibia of aged mice. Age (Dordr) 38, 38.

Ashpole, N.M., Herron, J.C., Mitschelen, M.C., Farley, J.A., Logan, S., Yan, H., Ungvari, Z., Hodges, E.L., Csiszar, A., Ikeno, Y., Humphrey, M.B., and Sonntag, W.E. (2015a). IGF-1 Regulates Vertebral Bone Aging Through Sex-Specific and Time-Dependent Mechanisms. J Bone Miner Res.

Ashpole, N.M., and Hudmon, A. (2011). Excitotoxic neuroprotection and vulnerability with CaMKII inhibition. Mol Cell Neurosci 46, 720–730.

Ashpole, N.M., Logan, S., Yabluchanskiy, A., Mitschelen, M.C., Yan, H., Farley, J.A., Hodges, E.L., Ungvari, Z., Csiszar, A., Chen, S., Georgescu, C., Hubbard, G.B., Ikeno, Y., and Sonntag, W.E. (2017). IGF-1 has sexually dimorphic, pleiotropic, and time-dependent effects on healthspan, pathology, and lifespan. Geroscience 39, 129–145.

Ashpole, N.M., Sanders, J.E., Hodges, E.L., Yan, H., and Sonntag, W.E. (2015b). Growth hormone, insulinlike growth factor-1 and the aging brain. Exp Gerontol 68, 76–81.

Bake, S., Okoreeh, A.K., Alaniz, R.C., and Sohrabji, F. (2016). Insulin-Like Growth Factor (IGF)-I Modulates Endothelial Blood-Brain Barrier Function in Ischemic Middle-Aged Female Rats. Endocrinology 157, 61–69.

Bake, S., Selvamani, A., Cherry, J., and Sohrabji, F. (2014). Blood brain barrier and neuroinflammation are critical targets of IGF-1-mediated neuroprotection in stroke for middle-aged female rats. PLoS One 9, e91427.

Benjamin, E.J., Muntner, P., Alonso, A., Bittencourt, M.S., Callaway, C.W., Carson, A.P., Chamberlain, A.M., Chang, A.R., Cheng, S., Das, S.R., Delling, F.N., Djousse, L., Elkind, M.S.V., Ferguson, J.F., Fornage, M., Jordan, L.C., Khan, S.S., Kissela, B.M., Knutson, K.L., Kwan, T.W., Lackland, D.T., Lewis, T.T., Lichtman, J.H., Longenecker, C.T., Loop, M.S., Lutsey, P.L., Martin, S.S., Matsushita, K., Moran, A.E., Mussolino, M.E., O’flaherty, M., Pandey, A., Perak, A.M., Rosamond, W.D., Roth, G.A., Sampson, U.K.A., Satou, G.M., Schroeder, E.B., Shah, S.H., Spartano, N.L., Stokes, A., Tirschwell, D.L., Tsao, C.W., Turakhia, M.P., Vanwagner, L.B., Wilkins, J.T., Wong, S.S., Virani, S.S., American Heart Association Council On, E., Prevention Statistics, C., and Stroke Statistics, S. (2019). Heart Disease and Stroke Statistics-2019 Update: A Report From the American Heart Association. Circulation 139, e56–e528.

Bradley, S.J., and Challiss, R.A. (2012). G protein-coupled receptor signalling in astrocytes in health and disease: a focus on metabotropic glutamate receptors. Biochem Pharmacol 84, 249–259.

Brown, L.S., Foster, C.G., Courtney, J.M., King, N.E., Howells, D.W., and Sutherland, B.A. (2019). Pericytes and Neurovascular Function in the Healthy and Diseased Brain. Front Cell Neurosci 13, 282.

Castilla-Cortazar, I., Aguirre, G.A., Femat-Roldan, G., Martin-Estal, I., and Espinosa, L. (2020). Is insulin-like growth factor-1 involved in Parkinson’s disease development? J Transl Med 18, 70.

Ceprian, M., and Fulton, D. (2019). Glial Cell AMPA Receptors in Nervous System Health, Injury and Disease. Int J Mol Sci 20.

Chen, W., He, B., Tong, W., Zeng, J., and Zheng, P. (2019). Astrocytic Insulin-Like Growth Factor-1 Protects Neurons Against Excitotoxicity. Front Cell Neurosci 13, 298.

Choi, D.W. (2020). Excitotoxicity: Still Hammering the Ischemic Brain in 2020. Front Neurosci 14, 579953.

Choi, D.W., Maulucci-Gedde, M., and Kriegstein, A.R. (1987). Glutamate neurotoxicity in cortical cell culture. J Neurosci 7, 357–368.

Choi, D.W., and Rothman, S.M. (1990). The role of glutamate neurotoxicity in hypoxic-ischemic neuronal death. Annu Rev Neurosci 13, 171–182.

De Smedt, A., Brouns, R., Uyttenboogaart, M., De Raedt, S., Moens, M., Wilczak, N., Luijckx, G.J., and De Keyser, J. (2011). Insulin-like growth factor I serum levels influence ischemic stroke outcome. Stroke 42, 2180–2185.

Farias Quipildor, G.E., Mao, K., Hu, Z., Novaj, A., Cui, M.H., Gulinello, M., Branch, C.A., Gubbi, S., Patel, K., Moellering, D.R., Tarantini, S., Kiss, T., Yabluchanskiy, A., Ungvari, Z., Sonntag, W.E., and Huffman, D.M. (2019). Central IGF-1 protects against features of cognitive and sensorimotor decline with aging in male mice. Geroscience 41, 185–208.

Fulop, G.A., Ramirez-Perez, F.I., Kiss, T., Tarantini, S., Valcarcel Ares, M.N., Toth, P., Yabluchanskiy, A., Conley, S.M., Ballabh, P., Martinez-Lemus, L.A., Ungvari, Z., and Csiszar, A. (2019). IGF-1 Deficiency Promotes Pathological Remodeling of Cerebral Arteries: A Potential Mechanism Contributing to the Pathogenesis of Intracerebral Hemorrhages in Aging. J Gerontol A Biol Sci Med Sci 74, 446–454.

George, P.M., and Steinberg, G.K. (2015). Novel Stroke Therapeutics: Unraveling Stroke Pathophysiology and Its Impact on Clinical Treatments. Neuron 87, 297–309.

Ghazi Sherbaf, F., Mohajer, B., Ashraf-Ganjouei, A., Mojtahed Zadeh, M., Javinani, A., Sanjari Moghaddam, H., Shirin Shandiz, M., and Aarabi, M.H. (2018). Serum Insulin-Like Growth Factor-1 in Parkinson’s Disease; Study of Cerebrospinal Fluid Biomarkers and White Matter Microstructure. Front Endocrinol (Lausanne) 9, 608.

Gubbi, S., Quipildor, G.F., Barzilai, N., Huffman, D.M., and Milman, S. (2018). 40 YEARS of IGF1: IGF1: the Jekyll and Hyde of the aging brain. J Mol Endocrinol 61, T171–T185.

Hayes, C.A., Valcarcel-Ares, M.N., and Ashpole, N.M. (2021). Preclinical and clinical evidence of IGF-1 as a prognostic marker and acute intervention with ischemic stroke. J Cereb Blood Flow Metab, 271678X211000894.

Higashi, Y., Sukhanov, S., Shai, S.Y., Danchuk, S., Snarski, P., Li, Z., Hou, X., Hamblin, M.H., Woods, T.C., Wang, M., Wang, D., Yu, H., Korthuis, R.J., Yoshida, T., and Delafontaine, P. (2020). Endothelial deficiency of insulin-like growth factor-1 receptor reduces endothelial barrier function and promotes atherosclerosis in Apoe-deficient mice. Am J Physiol Heart Circ Physiol 319, H730–H743.

Johnsen, S.P., Sorensen, H.T., Thomsen, J.L., Gronbaek, H., Flyvbjerg, A., Engberg, M., and Lauritzen, T. (2004). Markers of fetal growth and serum levels of insulin-like growth factor (IGF) I,-II and IGF binding protein 3 in adults. Eur J Epidemiol 19, 41–47.

Johnson, W., Onuma, O., Owolabi, M., and Sachdev, S. (2016). Stroke: a global response is needed. Bull World Health Organ 94, 634–634A.

Katzan, I.L., Furlan, A.J., Lloyd, L.E., Frank, J.I., Harper, D.L., Hinchey, J.A., Hammel, J.P., Qu, A., and Sila, C.A. (2000). Use of tissue-type plasminogen activator for acute ischemic stroke: the Cleveland area experience. JAMA 283, 1151–1158.

Khennouf, L., Gesslein, B., Brazhe, A., Octeau, J.C., Kutuzov, N., Khakh, B.S., and Lauritzen, M. (2018). Active role of capillary pericytes during stimulation-induced activity and spreading depolarization. Brain 141, 2032–2046.

Kisler, K., Nelson, A.R., Rege, S.V., Ramanathan, A., Wang, Y., Ahuja, A., Lazic, D., Tsai, P.S., Zhao, Z., Zhou, Y., Boas, D.A., Sakadzic, S., and Zlokovic, B.V. (2017). Pericyte degeneration leads to neurovascular uncoupling and limits oxygen supply to brain. Nat Neurosci 20, 406–416.

Kisler, K., Nikolakopoulou, A.M., Sweeney, M.D., Lazic, D., Zhao, Z., and Zlokovic, B.V. (2020). Acute Ablation of Cortical Pericytes Leads to Rapid Neurovascular Uncoupling. Front Cell Neurosci 14, 27.

Kleindorfer, D., Lindsell, C.J., Brass, L., Koroshetz, W., and Broderick, J.P. (2008). National US estimates of recombinant tissue plasminogen activator use: ICD-9 codes substantially underestimate. Stroke 39, 924–928.

Lalo, U., Pankratov, Y., Parpura, V., and Verkhratsky, A. (2011). Ionotropic receptors in neuronal-astroglial signalling: what is the role of “excitable” molecules in non-excitable cells. Biochim Biophys Acta 1813, 992–1002.

Li, Y., Sun, W., Han, S., Li, J., Ding, S., Wang, W., and Yin, Y. (2017). IGF-1-Involved Negative Feedback of NR2B NMDA Subunits Protects Cultured Hippocampal Neurons Against NMDA-Induced Excitotoxicity. Mol Neurobiol 54, 684–696.

Liu, X.F., Fawcett, J.R., Thorne, R.G., and Frey, W.H., 2nd (2001). Non-invasive intranasal insulin-like growth factor-I reduces infarct volume and improves neurologic function in rats following middle cerebral artery occlusion. Neurosci Lett 308, 91–94.

Lopez-Lopez, C., Leroith, D., and Torres-Aleman, I. (2004). Insulin-like growth factor I is required for vessel remodeling in the adult brain. Proc Natl Acad Sci U S A 101, 9833–9838.

Moskowitz, M.A., Lo, E.H., and Iadecola, C. (2010). The science of stroke: mechanisms in search of treatments. Neuron 67, 181–198.

National Institute of Neurological, D., and Stroke Rt, P.a.S.S.G. (1995). Tissue plasminogen activator for acute ischemic stroke. N Engl J Med 333, 1581–1587.

Parker, K., Berretta, A., Saenger, S., Sivaramakrishnan, M., Shirley, S.A., Metzger, F., and Clarkson, A.N. (2017). PEGylated insulin-like growth factor-I affords protection and facilitates recovery of lost functions post-focal ischemia. Sci Rep 7, 241.

Prabhu, D., Khan, S.M., Blackburn, K., Marshall, J.P., and Ashpole, N.M. (2019). Loss of insulin-like growth factor-1 signaling in astrocytes disrupts glutamate handling. J Neurochem.

Qureshi, A.I., Suri, M.F., Nasar, A., He, W., Kirmani, J.F., Divani, A.A., Prestigiacomo, C.J., and Low, R.B. (2005). Thrombolysis for ischemic stroke in the United States: data from National Hospital Discharge Survey 1999-2001. Neurosurgery 57, 647–654; discussion 647-654.

Rosenberg, P.A., Amin, S., and Leitner, M. (1992). Glutamate uptake disguises neurotoxic potency of glutamate agonists in cerebral cortex in dissociated cell culture. J Neurosci 12, 56–61.

Saber, H., Himali, J.J., Beiser, A.S., Shoamanesh, A., Pikula, A., Roubenoff, R., Romero, J.R., Kase, C.S., Vasan, R.S., and Seshadri, S. (2017). Serum Insulin-Like Growth Factor 1 and the Risk of Ischemic Stroke: The Framingham Study. Stroke 48, 1760–1765.

Serhan, A., Boddeke, E., and Kooijman, R. (2019). Insulin-Like Growth Factor-1 Is Neuroprotective in Aged Rats With Ischemic Stroke. Front Aging Neurosci 11, 349.

Serrano, A., Robitaille, R., and Lacaille, J.C. (2008). Differential NMDA-dependent activation of glial cells in mouse hippocampus. Glia 56, 1648–1663.

Sonntag, W.E., Deak, F., Ashpole, N., Toth, P., Csiszar, A., Freeman, W., and Ungvari, Z. (2013). Insulin-like growth factor-1 in CNS and cerebrovascular aging. Front Aging Neurosci 5, 27.

Stanimirovic, D.B., and Friedman, A. (2012). Pathophysiology of the neurovascular unit: disease cause or consequence? J Cereb Blood Flow Metab 32, 1207–1221.

Sun, J., Huang, Y., Gong, J., Wang, J., Fan, Y., Cai, J., Wang, Y., Qiu, Y., Wei, Y., Xiong, C., Chen, J., Wang, B., Ma, Y., Huang, L., Chen, X., Zheng, S., Huang, W., Ke, Q., Wang, T., Li, X., Zhang, W., Xiang, A.P., and Li, W. (2020). Transplantation of hPSC-derived pericyte-like cells promotes functional recovery in ischemic stroke mice. Nat Commun 11, 5196.

Tang, J.H., Ma, L.L., Yu, T.X., Zheng, J., Zhang, H.J., Liang, H., and Shao, P. (2014). Insulin-like growth factor-1 as a prognostic marker in patients with acute ischemic stroke. PLoS One 9, e99186.

Tarantini, S., Balasubramanian, P., Yabluchanskiy, A., Ashpole, N.M., Logan, S., Kiss, T., Ungvari, A., Nyul-Toth, A., Schwartzman, M.L., Benyo, Z., Sonntag, W.E., Csiszar, A., and Ungvari, Z. (2021). IGF1R signaling regulates astrocyte-mediated neurovascular coupling in mice: implications for brain aging. Geroscience 43, 901–911.

Tarantini, S., Giles, C.B., Wren, J.D., Ashpole, N.M., Valcarcel-Ares, M.N., Wei, J.Y., Sonntag, W.E., Ungvari, Z., and Csiszar, A. (2016a). IGF-1 deficiency in a critical period early in life influences the vascular aging phenotype in mice by altering miRNA-mediated post-transcriptional gene regulation: implications for the developmental origins of health and disease hypothesis. Age (Dordr) 38, 239–258.

Tarantini, S., Tucsek, Z., Valcarcel-Ares, M.N., Toth, P., Gautam, T., Giles, C.B., Ballabh, P., Wei, J.Y., Wren, J.D., Ashpole, N.M., Sonntag, W.E., Ungvari, Z., and Csiszar, A. (2016b). Circulating IGF-1 deficiency exacerbates hypertension-induced microvascular rarefaction in the mouse hippocampus and retrosplenial cortex: implications for cerebromicrovascular and brain aging. Age (Dordr) 38, 273–289.

Tarantini, S., Valcarcel-Ares, N.M., Yabluchanskiy, A., Springo, Z., Fulop, G.A., Ashpole, N., Gautam, T., Giles, C.B., Wren, J.D., Sonntag, W.E., Csiszar, A., and Ungvari, Z. (2017). Insulin-like growth factor 1 deficiency exacerbates hypertension-induced cerebral microhemorrhages in mice, mimicking the aging phenotype. Aging Cell 16, 469–479.

Terasaki, Y., Liu, Y., Hayakawa, K., Pham, L.D., Lo, E.H., Ji, X., and Arai, K. (2014). Mechanisms of neurovascular dysfunction in acute ischemic brain. Curr Med Chem 21, 2035–2042.

Toth, P., Tarantini, S., Ashpole, N.M., Tucsek, Z., Milne, G.L., Valcarcel-Ares, N.M., Menyhart, A., Farkas, E., Sonntag, W.E., Csiszar, A., and Ungvari, Z. (2015a). IGF-1 deficiency impairs neurovascular coupling in mice: implications for cerebromicrovascular aging. Aging Cell 14, 1034–1044.

Toth, P., Tarantini, S., Ashpole, N.M., Tucsek, Z., Milne, G.L., Valcarcel-Ares, N.M., Menyhart, A., Farkas, E., Sonntag, W.E., Csiszar, A., and Ungvari, Z. (2015b). IGF-1 deficiency impairs neurovascular coupling in mice: implications for cerebromicrovascular aging. Aging Cell.

Toth, P., Tarantini, S., Csiszar, A., and Ungvari, Z. (2017). Functional vascular contributions to cognitive impairment and dementia: mechanisms and consequences of cerebral autoregulatory dysfunction, endothelial impairment, and neurovascular uncoupling in aging. Am J Physiol Heart Circ Physiol 312, H1–H20.

Wang, Y., Wang, W., Li, D., Li, M., Wang, P., Wen, J., Liang, M., Su, B., and Yin, Y. (2014). IGF-1 alleviates NMDA-induced excitotoxicity in cultured hippocampal neurons against autophagy via the NR2B/PI3K-AKT-mTOR pathway. J Cell Physiol 229, 1618–1629.

Watanabe, T., Miyazaki, A., Katagiri, T., Yamamoto, H., Idei, T., and Iguchi, T. (2005). Relationship between serum insulin-like growth factor-1 levels and Alzheimer’s disease and vascular dementia. J Am Geriatr Soc 53, 1748–1753.

Yang, S., Jin, H., Zhu, Y., Wan, Y., Opoku, E.N., Zhu, L., and Hu, B. (2017). Diverse Functions and Mechanisms of Pericytes in Ischemic Stroke. Curr Neuropharmacol 15, 892–905.

